# Enhancing reproducibility and decentralization in single cell research with biocytometry

**DOI:** 10.1101/2024.07.01.601489

**Authors:** Pavel Fikar, Laura Alvarez, Laura Berne, Martin Cienciala, Christopher Kan, Hynek Kasl, Mona Luo, Zuzana Novackova, Sheyla Ordonez, Zuzana Sramkova, Monika Holubova, Daniel Lysak, Lyndsay Avery, Andres A. Caro, Roslyn N. Crowder, Laura A. Diaz-Martinez, David W. Donley, Rebecca R. Giorno, Irene K. Guttilla Reed, Lori L. Hensley, Kristen C. Johnson, Paul Kim, Audrey Y. Kim, Adriana J. LaGier, Jamie J. Newman, Elizabeth Padilla-Crespo, Nathan S. Reyna, Nikolaos Tsotakos, Noha N. Al-Saadi, Tayler Appleton, Ana Arosemena-Pickett, Braden A. Bell, Grace Bing, Bre Bishop, Christa Forde, Michael J. Foster, Kassidy Gray, Bennett L. Hasley, Kennedy Johnson, Destiny Jen’a Jones, Allison C. LaShall, Kennedy McGuire, Naomi McNaughton, Angelina M. Morgan, Lucas Norris, Landon A. Ossman, Paollette A. Rivera-Torres, Madeline E. Robison, Kathryn Thibodaux, Lescia Valmond, Daniel Georgiev

**Affiliations:** Department of Research & Development, Sampling Human Inc., Berkeley, California, 94720, USA; Department of Hematology and Oncology, University Hospital Pilsen, Pilsen, 30100, Czech Republic; Department of Biology, Saint Michael’s College, Colchester, Vermont, 05439, USA; Chemistry Department, Hendrix College, Conway, Arkansas, 72032, USA; Department of Biology, Stetson University, Deland, Florida, 32723, USA; Department of Biology, Gonzaga University, Spokane, Washington, 99258, USA; Department of Biology, Harding University, Searcy, Arkansas, 72143, USA; School of Biological Sciences, Louisiana Tech University, Ruston, Louisiana, 71272, USA; Department of Biology, University of Saint Joseph, West Hartford, Connecticut, 06117, USA; Department of Biological Sciences, Jacksonville State University, Jacksonville, Alabama, 36265, USA; Department of Life Sciences, University of New Hampshire, Manchester, New Hampshire, 03101, USA; Department of Biological Sciences, Grambling State University, Grambling, Louisiana, 71245, USA; Department of Biology, College of Social and Natural Sciences, Grand View University, Des Moines, Iowa, 50316, USA; Department of Science and Technology, Inter American University of Puerto Rico-Aguadilla, Aguadilla, Puerto Rico, 00605, USA; Department of Biology, Ouachita Baptist University, Arkadelphia, Arkansas, 71929, USA; Department of Biological Sciences, School of Science Engineering and Technology, Penn State Harrisburg, Middletown, Pennsylvania, 17057, USA; Faculty of Applied Sciences, University of West Bohemia, Pilsen, 30100, Czech Republic

## Abstract

Biomedicine today is experiencing a shift towards decentralized data collection, which promises enhanced reproducibility and collaboration across diverse laboratory environments. This inter-laboratory study evaluates the performance of biocytometry, a method utilizing engineered bioparticles for enumerating cells based on their surface antigen patterns. In a decentralized framework, spanning 78 assays conducted by 30 users across 12 distinct laboratories, biocytometry consistently demonstrated significant statistical power in discriminating numbers of target cells at varying concentrations as low as 1 cell per 100,000 background cells. User skill levels varied from expert to beginner capturing a range of proficiencies. Measurement was performed in a decentralized environment without any instrument cross-calibration or advanced user training outside of a basic instruction manual. The results affirm biocytometry to be a viable solution for immunophenotyping applications demanding sensitivity as well as scalability and reproducibility and paves the way for decentralized analysis of rare cells in heterogeneous samples.

## Introduction

Reproducibility, long held as the gold standard in scientific research, is now being critically examined, facing questions and challenges from the broader research community [1]. Single cell research is at the forefront of the reproducibility challenge, facing significant inter-laboratory variability [2–4] and the sobering observation that a mere 11% of preclinical studies are successfully reproduced [5]. Flow cytometry, a cornerstone in clinical single cell analysis, continues to pose reproducibility problems. Despite concerted efforts by consortia like EuroFlow, The ONE Study, and HIPC to standardize its methodologies [6], challenges persist, particularly in the quantification of rare cells which exhibit pronounced variability [7]. Yet, the analysis of cell populations with low frequencies is indispensable across the entire spectrum of care.

Sensitive single-cell data is highly impactful, from the early stages of disease detection and progression monitoring [8–10] to the evaluation of therapeutic interventions [11] and the surveillance of residual disease [12]. Rare cell populations are often crucial to treating or diagnosing disease, such as Parkinson’s associated with certain microglial populations or immunosuppression instigated by regulatory T cell activity. Detection and enumeration of rare cells, however, remains a challenge even for expert clinical laboratories due to complexity of the underlying measurement technologies as well as the unavoidable heterogeneity in the cell population and sample matrix caused by natural and induced phenomena. These limitations have hindered scalability of medical research and the application of single cell data in medical practice. For example, most current flow cytometry assays are lab-developed tests (LDTs) that are designed, manufactured and validated within a single laboratory. This is in stark contrast to molecular diagnostics that can be scaled to 100s of millions of tests globally. New technologies are necessary to increase reproducibility and adoption of cellular data in medicine.

This investigation centers on reproducibility of biocytometry, a method that utilizes engineered bioparticles for target cell identification [13]. Inspired by natural cellular identification mechanisms, biocytometry allows for simultaneous identification of target cell immunophenotypes in suspension. The system employs bioparticles engineered to report the presence of specific surface antigen patterns. When introduced to a sample, the bioparticles bind to surface antigens present on cells in suspension. As they bind, the bioparticles react to adjacency of other bioparticles with a strong release of a luminescent reporter into the medium, facilitating sensitive enumeration of target cells. This approach has been demonstrated to support enumeration of various immunophenotypes defined by known combinations of surface antigens, e.g., T-cell activation, cell type specific apoptosis, cancer cell biomarkers. Unlike traditional analytical approaches, it overcomes the constraints tied to use of instruments and molecular probes, introducing a streamlined workflow that bodes well for improved reproducibility, especially in high-sensitivity applications.

To evaluate the reproducibility of biocytometry, an extensive interlaboratory study was initiated in collaboration with the Cell Biology Education Consortium (CBEC), aiming to assess performance in high sensitivity sample analysis in both centralized and decentralized data collection modes. Participants spanned a spectrum from novice undergraduate students to experienced professionals, ensuring comprehensive evaluation across varying proficiency levels. Evaluation was performed on pre-formulated human mockup (HUMO) samples comprising target cells (HaCaT cell line, EpCAM+) integrated into the leukocyte surrogate population (HL-60 cell line, EpCAM-). Target cells were present at varying concentrations simulating different sample types - negative (absent of target cells), low (10^-5^ sensitivity equivalent), and high (10^-4^ sensitivity equivalent), providing a robust framework for assessing the technology’s precision across different sensitivity levels.

The findings of this study position biocytometry as a significant advancement over current cellular analysis methods, given its marked reproducibility and sensitivity. Our analysis draws upon a dataset derived from 78 assays, undertaken by 30 participants across 12 distinct laboratories. The initial observations indicate that biocytometry differentiates between analyzed HUMO samples with notable statistical significance (p = 3×10^-5^), a finding further affirmed by cross-referencing absolute target cell counts via fluorescence microscopy. The assay demonstrated remarkable user versatility, as results remained consistent regardless of the proficiency spectrum, encompassing even non-trained participants. Crucially, both centralized and decentralized data collection modes yielded equivalent statistical power. Adoption of biocytometry is expected to bolster collection of single-cell data in applications akin to decentralized clinical trials, allowing for comprehensive and statistically significant analysis across diverse research environments.

## Results

### Study

To evaluate the sensitivity, accuracy, and reproducibility of biocytometry across diverse settings, we organized an interlaboratory study in collaboration with participants from multiple institutions (78 assays, 30 participants, 12 laboratories). This diverse group encompassed proficient users experienced in biocytometry and external participants new to this technology. It included seasoned principal investigators as well as undergraduate students, each bringing different levels of laboratory expertise. This varied proficiency among participants enabled a detailed and comprehensive evaluation of biocytometry’s applicability and robustness across different user groups, while the involvement of multiple institutions was instrumental in revealing how biocytometry performs across a range of laboratory settings, each equipped with different instruments, thereby providing a thorough understanding of its performance in diverse environments.

Every participant was supplied a biocytometry kit for the detection and enumeration of EpCAM+ target cells. Each kit contained all necessary consumables and reagents required for the biocytometry protocol (see Fig 1A). Along with the kit, we provided a set of synthetic human mockup (HUMO) samples containing various concentrations of EpCAM+ cells for analysis (see Fig 1B). A notable aspect of this study was the reliance on standard laboratory equipment already present in most research settings, such as pipettes, microcentrifuges, thermocyclers, and luminescence readers. To facilitate the decentralized data collection, the biocytometry kits and HUMO samples were first shipped from the centralized laboratory to the reference laboratory and then distributed to the designated target laboratories (see Fig 1C).

**Fig 1.**
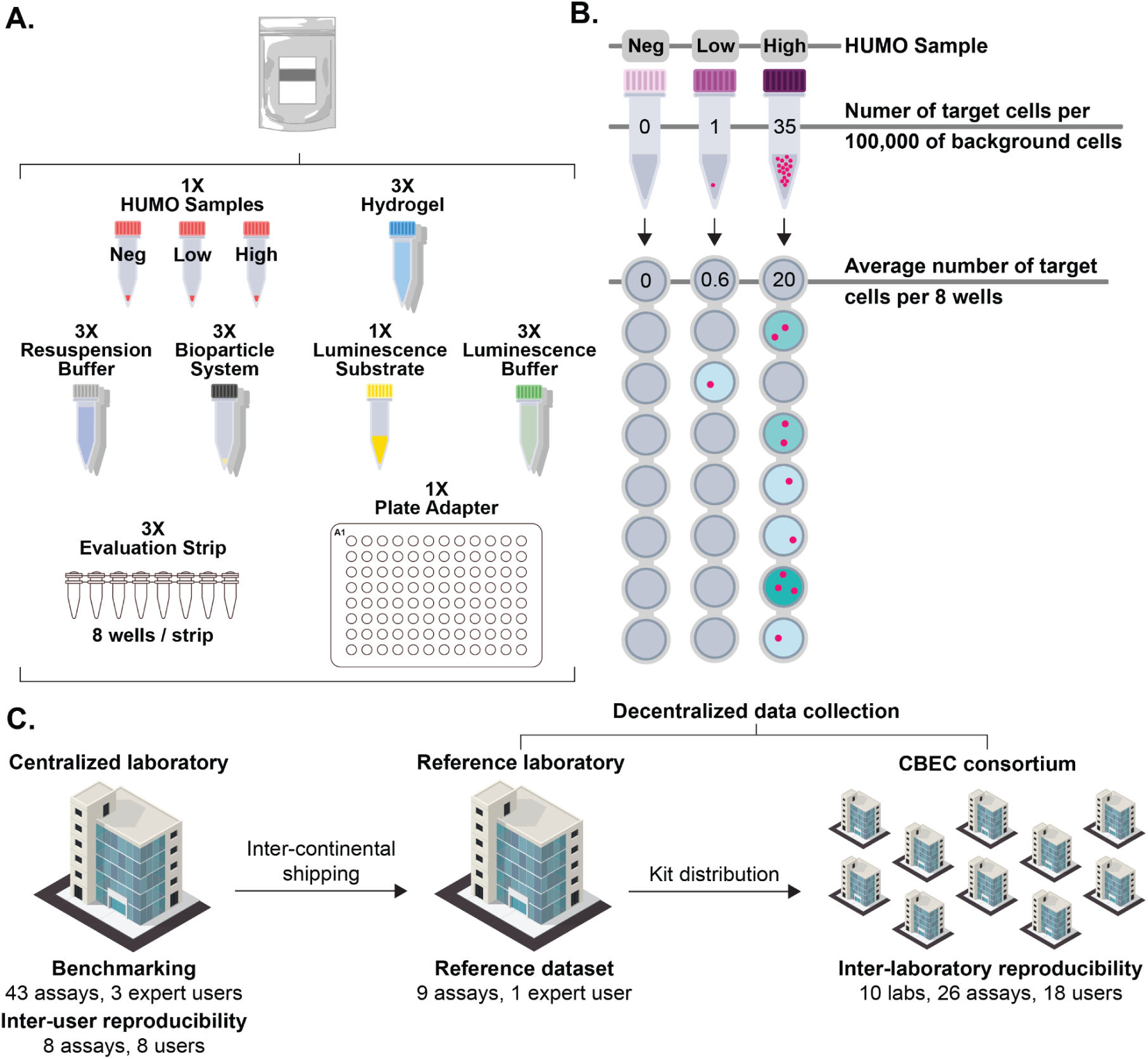
Study design. **A)** Each kit includes all consumables and reagents required for the biocytometry protocol. These comprise the bioparticle reagent, a set of HUMO samples (negative, low and high), a hydrogel-based incubation medium, resuspension and luminescence buffers, a luminescence substrate, evaluation strips and a plate adapter for the final luminescence readout in a microplate reader. **B)** HUMO samples of 100,000 cells were provided with each biocytometry kit. The concentrations of targets in the three HUMO samples were: 0 in 100,000 cells (negative), 1 in 100,000 cells (low), and 35 in 100,000 cells (high). For measurement, 56,000 cells were analyzed across 8 wells. The figure illustrates how signal-to-noise ratio values and the target count estimates are proportional to, and derived from, the luminescence readout of individual sample wells. **C)** Distribution of all 78 assays from the study dataset, each encompassing HUMO samples of every type (negative, low and high). The figure outlines the distribution of the biocytometry kits and HUMO samples and the number of assays carried out by various laboratories.

The study was structured in three stages dedicated to i) biocytometry benchmarking, ii) inter-user reproducibility and iii) decentralized reproducibility. In benchmarking biocytometry, we validated the sensitivity, accuracy, and reproducibility of the method on the given samples in a centralized and controlled setting. This stage was carried out by three proficient users and established a basis for the method’s capabilities under ideal conditions.

The inter-user reproducibility study included 8 users of various proficiencies performing specific assays in a centralized location, utilizing the same instrumentation. Potential hardware discrepancies were eliminated, thereby allowing us to isolate the impact of user idiosyncrasies. Potential sources of variability specific to the biocytometry protocol include hydrogel setting, overall timing, handling vigor, and exposure of light-sensitive reagents.

The decentralized reproducibility study enlisted external participants from multiple institutions to conduct biocytometry assays in various laboratory settings. This stage was instrumental in validating the consistency and reliability of biocytometry in real-world, decentralized conditions. All participants were provided with standardized instruction materials. This included a written protocol and a 10-minute instructional video tutorial. Participants conducted assays using their own laboratory equipment and were provided no real-time or hands-on guidance.

### Samples

Three types of HUMO samples were prepared for benchmarking, and the inter-user and decentralized reproducibility studies. One of each HUMO sample type was included in each biocytometry kit distributed to participants. These samples were composed of varying numbers of HaCaT (keratinocyte) EpCAM+ cells, designated as the target cells, and a consistent number of approximately 100,000 background cells derived from HL-60 (promyelocytic) EpCAM- cell line, representing human leukocytes. The selection of EpCAM+ cells as the target in this study provides a relevant and practical model that illustrates the potential of biocytometry in critical areas like cancer diagnostics and single cell research and its capability to handle clinically relevant biomarkers.

The three types of HUMO samples varied in their proportions of target cells (see Fig 1B). The target cell concentrations were chosen to demonstrate the capabilities of biocytometry in “rare-event analysis”, typically defined at frequencies of 1 in 1000 or below [8]. Negative HUMO samples contained no target cells to establish the method’s false positive rate. This is critical as high noise levels in techniques like flow cytometry can make it statistically challenging to distinguish between samples with no targets and those with a low number of targets. The low HUMO sample, representing a 10^-5^ equivalent, contained 1 target cell per 100,000 background cells on average. It was specifically designed to emulate a scenario with a single target cell within a large population of background cells. The high HUMO sample, representing a 10^-4^ equivalent, contained 35 target cells per 100,000 background cells on average, illustrating the wider sensitivity range of biocytometry. To preserve the integrity of the blind study, each sample was labeled with a disguised nomenclature, effectively concealing its true identity.

### Biocytometric analysis

The biocytometry protocol includes three basic steps: sample and bioparticle mixing, incubation, and luminescence measurement. Each assay consumed approximately 30 minutes of hands-on time and was complete in roughly 5 hours total time. When performed in batches, up to 12 samples could be processed together. For detailed protocol see Materials and methods.

Participants submitted the results in the form of averaged luminescence values online without the need for additional adjustments. In addition, metadata regarding the plate reader model, settings, and additional comments were collected. Participants were asked to self-report any deviations from the protocol. While assays significantly deviating from the protocol were excluded, no data-based exclusion criteria were employed. Assays performed using readers with no or too low sensitivity to luminescence were excluded from further analysis.

In the analysis of a biocytometry assay, five key values were computed from the raw luminescence data: SNR_WELL_, SNR_SAMPLE_, T_WELL_, T_SAMPLE_ and SNRT (see Materials and Methods). SNR_WELL_ is the signal-to-noise ratio (SNR) calculated for each well. SNR_SAMPLE_ is the sum of the SNR_WELL_ values calculated for each sample. The number of target cells per well is denoted by T_WELL_ and is equal to SNR_WELL_ normalized by the target-specific conversion factor CF evaluated during benchmarking of biocytometry. The number of target cells per sample T_SAMPLE_ equals the sum of T_WELL_ for each sample. Lastly, for benchmarking purposes, select measurements were paired with microscopic images to compute the true signal-to-noise ratio per target SNRT. Estimates of this ratio are denoted as SNRT_EST_ and serve as further indicators of assay consistency in the absence of microscopy measurements.

Data were systematically categorized into two principal groups: centralized data, focusing on benchmarking of biocytometry and inter-user reproducibility; and decentralized data, emphasizing real-world application variability. Each assay was further characterized by additional metadata, including self-reported user proficiency level, which played a significant role in subsequent analyses, aiding in the comprehensive evaluation of biocytometry’s applicability and robustness.

### Benchmarking of biocytometry

HUMO samples were analyzed in 43 biocytometry assays carried out by 3 proficient users in a centralized laboratory. Target cell counts were obtained by analyzing the same samples using fluorescence microscopy, which served as a comparative standard. Target cells were fluorescently labeled during the sample preparation phase to enable the reliable differentiation of target cells from background cells during microscopy analysis. The microscopy-derived counts, in conjunction with the SNR values, facilitated the identification of the conversion factor (CF). Once established, this CF was utilized throughout the study to convert the SNR values into corresponding target estimates, T_WELL_ and T_SAMPLE_. Statistical analysis was subsequently performed, focusing on sample discrimination and reproducibility.

The CF value was determined through linear regression using the least squares method, as depicted in Fig 2A. This analysis reveals that the detection of one target cell by biocytometry corresponds to an approximate increase of 25.4 in the SNR for the corresponding well. The linear relationship between the SNR values and the number of target cells is clearly demonstrated, indicating that SNR values are directly proportional to the target cell count. This data highlights the random distribution of target cells within the sample volume and underscores the predictive power of biocytometry in accurately reflecting the true distribution of cells across a range of target cell concentrations.

**Fig 2.**
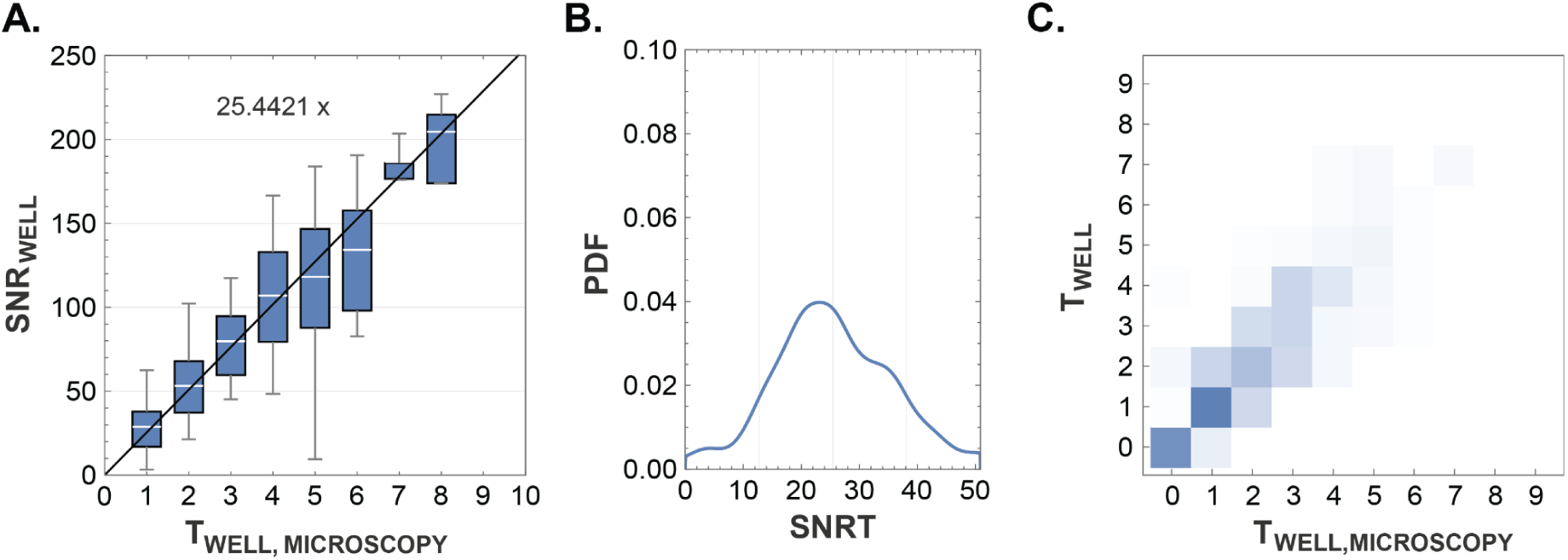
Validation of biocytometry using microscopy. **A)** A comparison of the SNR_WELL_ values reported by biocytometry and the target cell counts obtained by microscopic examination. The agreement validates the homogeneity in sample preparation and emphasizes the predictive power of biocytometry in capturing the true underlying distribution of cells. A conversion factor CF ∼ 25.4421 was identified by linear regression based with the least square method. **B)** A histogram visualizing the SNRT distribution derived from 302 HUMO sample wells evaluated by combination of biocytometry and microscopy. The alignment between the distribution (SNRT = 26.9, CV = 49.1%) and the nominal SNRT value (CF = 25.4) illustrates the reliability of biocytometry to enumerate target cells. **C)** Biocytometry vs microscopy confusion matrix illustrating a two-dimensional probability mass function (PMF). Higher probabilities were accentuated with more intense colors. The strong concordance between target estimates garnered via biocytometry and those obtained through microscopy underscores the reliable performance of biocytometry in estimating the number of target cells present in the sample. Any minor underestimation noted in microscopy can potentially be attributed to loss of target cells during the microscopy validation process.

SNRT values were evaluated for 302 HUMO sample wells from 43 biocytometry assays (wells containing no target cells omitted). Fig 4B presents the SNRT distribution (mean SNRT = 26.9, CV = 49.1%) in the form of a probability density function (PDF). The measured distribution is well aligned with the CF value of 25.4.

The target count estimates obtained by biocytometry were cross-validated with absolute target counts obtained by microscopy (see Fig 2C). The strong concordance between the two underscores the reliable performance of biocytometry in estimating the number of target cells present in the sample. Any minor underestimation noted in microscopy can potentially be attributed to loss of target cells during material transfer for the microscopy validation process.

Using the Wilcoxon signed-rank test, the mean SNR of the HUMO samples were found to be statistically distinct from one another: between negative and low HUMO (p = 3×10^-5^), between negative and high HUMO (p = 1×10^-8^), and between low and high HUMO (p = 1×10^-8^). These notably low p-values are particularly significant considering the minuscule differences in cellular composition between the negative and low HUMO samples, which only differ by a mere 0.001%. This demonstrates the remarkable sensitivity of biocytometry, as it can effectively discriminate closely related HUMO samples with high statistical significance (Fig 3A).

**Fig 3.**
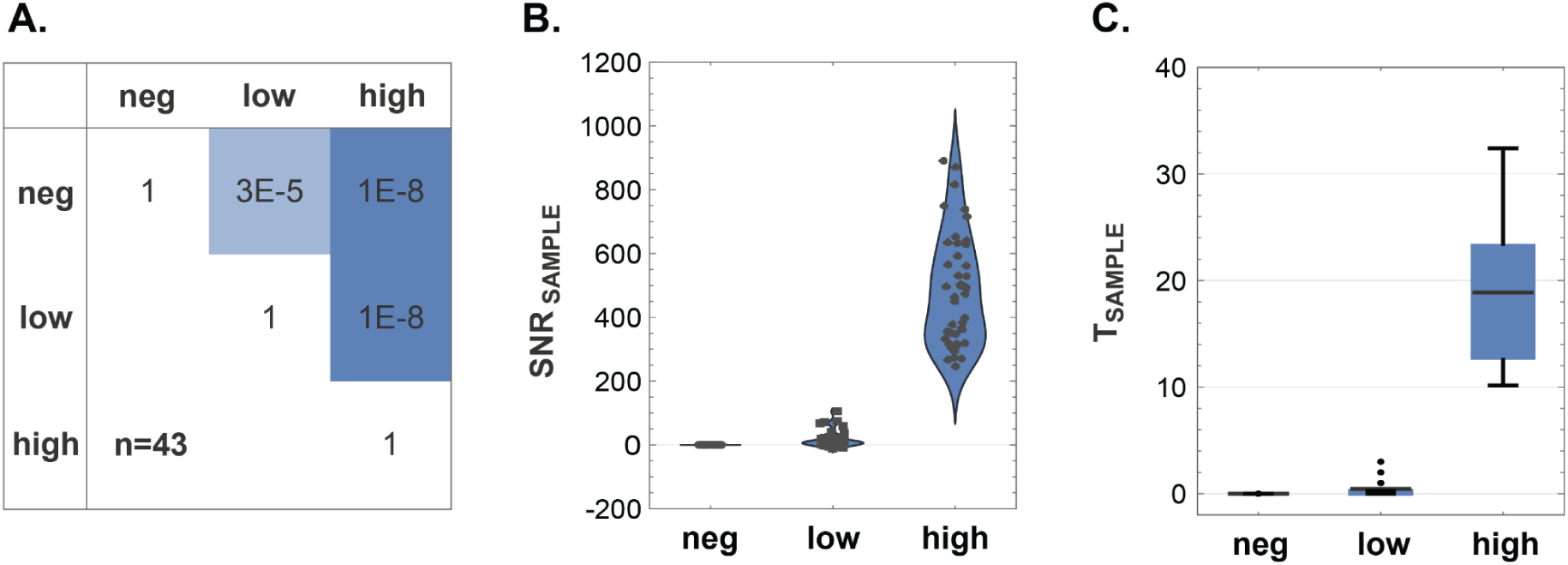
Discriminatory power of biocytometry. **A)** A confusion matrix showcasing the p-values calculated from the Wilcoxon signed-rank test demonstrates the ability of biocytometry to differentiate between different HUMO samples with statistical significance. **B)** Violin charts representing the distribution of SNR_SAMPLE_ values emphasize consistent performance of biocytometry across different HUMO samples. Average SNR values of 0.0, 15.6 and 483.6 were observed for the negative, low and high HUMO samples respectively. **C)** Target prediction of biocytometry is illustrated by T_SAMPLE_ values for each HUMO sample type. Average T_SAMPLE_ values of 0.0, 0.4 and 18.9 were observed for the negative, low and high HUMO samples respectively. Key statistical values, including means, quartiles and 5% and 95% quantiles, are represented within the box-and-whisker plots.

The discriminatory capacity is further highlighted by the distinct SNR_SAMPLE_ distributions depicted in Fig 3B and the corresponding T_SAMPLE_ values illustrated in Fig 3C. For the negative HUMO samples, an average SNR_SAMPLE_ of 0.0 was recorded, with a maximum absolute deviation from this average being 0.2. Such a narrow range of SNR values corroborates the absence of target cells with no false positives, signifying a precise specificity in the biocytometry measurements. For the low HUMO, across all assays, an average SNR_SAMPLE_ of 15.6 was measured. The number of targets for low HUMO was estimated to 0.4 target cells. This is in alignment with the expected cell concentration in the 10^-5^ sensitivity equivalent (see Fig 1B) and approaches the theoretical detection limit of a single target cell. In high HUMO samples, the elevated average SNR_SAMPLE_ of 483.6 was observed and the number of targets was estimated to 18.9 target cells in the analyzed sample volume. These results are in strong agreement with the expected 20 cells in the 10^-4^ sensitivity equivalent (see Fig 1C).

### Inter-user reproducibility study

The inter-user reproducibility study characterized consistency of biocytometry results across users with varying proficiency levels. This analysis encompasses the results of 51 assays: 43 assays performed by 3 users, complemented by 8 additional assays carried out by 8 unique users. All assays were executed utilizing the same instrumentation in a centralized laboratory.

The reproducibility across user proficiency levels is illustrated by consistent SNRT values evaluated across 137 wells by a combination of biocytometry and microscopy (Fig 4A). Routine, regular, and first-time users demonstrated average SNRT_WELL_ values of 18.9 (CV = 48.7%, 46 wells), 18.3 (CV = 60.6%, 23 wells), and 19.6 (CV = 53.1%, 68 wells), respectively, culminating in an aggregate mean of 19.2 (CV = 52.6%). From a mechanistic perspective, the alignment of the results shows the intrinsic mechanisms of biocytometry are such that they deliver consistent results irrespective of the proficiency of the user.

**Fig 4.**
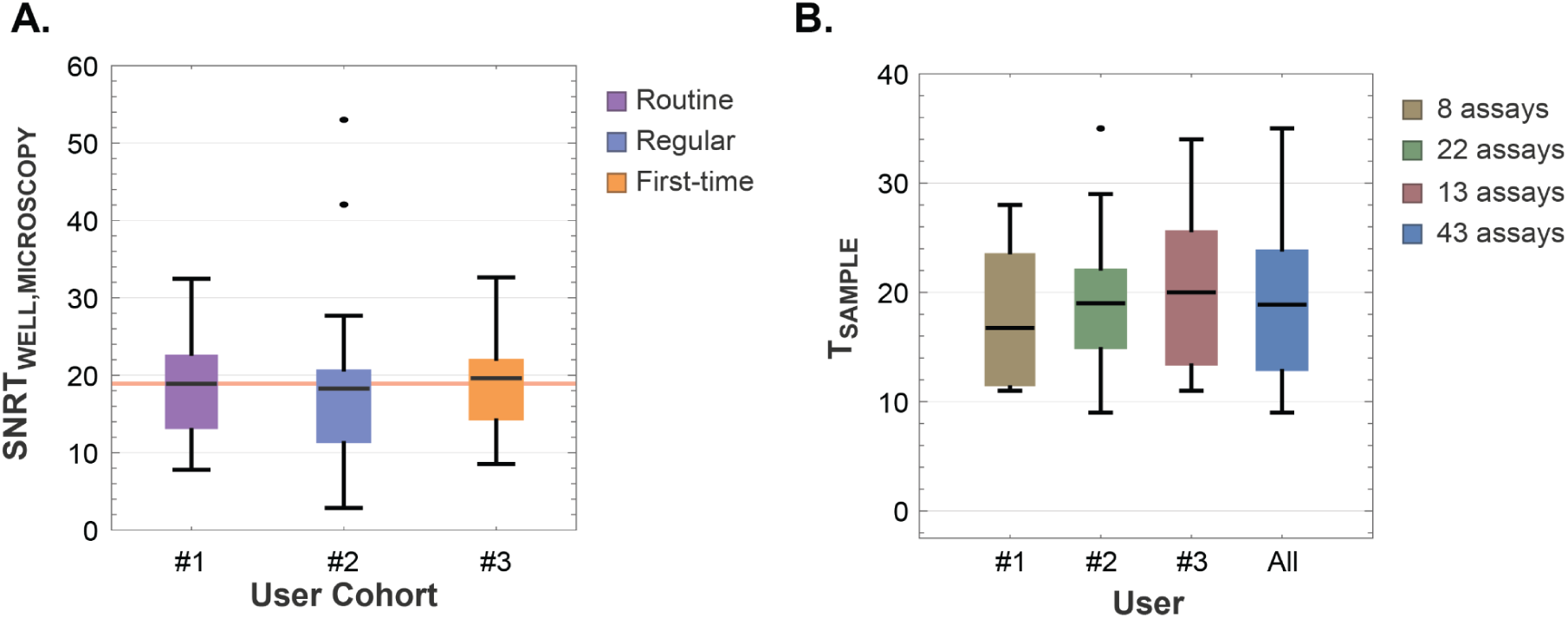
Assessment of inter-user reproducibility: A comparative study on SNRT values and *high HUMO* target estimates across different users. **A)** Distribution of SNRT_WELL_ values obtained for 137 wells by a combination of biocytometry and microscopy across users with varying proficiency levels. Average SNRT_WELL_ values of 18.9 (CV = 48.7%), 18.3 (CV = 60.6%), 19.6 (CV = 53.1%) were registered by the routine, regular and first-time users respectively. The consolidated mean of 19.2 (CV = 52.6%) is denoted by the red line. Key statistical values, including means, quartiles and 5% and 95% quantiles, are represented within the box-and-whisker plots. **B)** A comparative analysis of high HUMO sample target estimates across users confirms the consistency of biocytometry, regardless of the user performing the assay. The means are 16.8, 19.0, 20.0 and 18.9 respectively. Key statistical values, with means, quartiles, 5% and 95% quantiles and outliers are clearly indicated.

The uniformity of outcomes across various users was further supported by conducting a comparative evaluation of target estimates for high HUMO samples. Assays carried out by the same individual were grouped together for this analysis (Fig 4B). Means recorded by the three users were 16.8 (CV = 44.2%), 19.0 (CV = 33.1%), 20.0 (CV = 39.7%). The consolidated mean stood at 18.9 (CV = 36.7%). The higher CVs are anticipated due to the random distribution of target cells in samples. The salient feature is rapid convergence of results, demonstrating that biocytometry delivers consistent performance irrespective of user-to-user variability. This encompasses everything from the initial setup and manipulation of samples through to data interpretation.

Statistical robustness of these findings is reinforced by one-way analysis of variance (ANOVA), with a p-value of 0.841 for the SNRT_WELL_ values across users, indicating no statistically significant difference due to user variability (Fig 4A). Similarly, the target estimates for high HUMO samples yielded a p-value of 0.588 (Fig 4B). These p-values underscore the robustness and consistency of biocytometry, demonstrating its reliability and suitability for application in a variety of laboratory settings by users of differing experience levels. Hence, biocytometry is validated as a reproducible method, from sample preparation to data interpretation, across user cohorts.

### Decentralized reproducibility study

The decentralized study included 35 assays conducted by 19 users in 11 laboratories. Participants had no prior experience with biocytometry and varying amounts of lab experience. The involved laboratories spanned varying logistical conditions and settings. The ultimate goal of these experiments was to validate that, irrespective of whether data collection is undertaken in a centralized or decentralized manner, biocytometry maintains consistent statistical significance in discriminating HUMO samples. The discriminatory ability and mechanistic performance of biocytometry was further tested following international shipping by an expert user at a reference laboratory. This data also provided a benchmark for comparing results across labs.

P-values obtained through the Wilcoxon signed-rank test were used to compare mean SNR values across different HUMO samples and settings (Fig 5A). In the reference laboratory (n=9), we observed a p-value of 0.34 between negative and low HUMO, 9×10^-3^ between negative and high HUMO, and 9×10^-3^ between low and high HUMO. In the inter-laboratory data collection performed by the external participants (n=26), we observed a p-value of 1×10^-2^ between negative and low HUMO, 9×10^-6^ between negative and high HUMO, and 2×10^-5^ between low and high HUMO. Isolating data obtained only by undergraduate users (n=10), we observed a p-value of 0.61 between negative and low HUMO, 6×10^-3^ between negative and high HUMO, and 6×10^-3^ between low and high HUMO.

**Fig 5.**
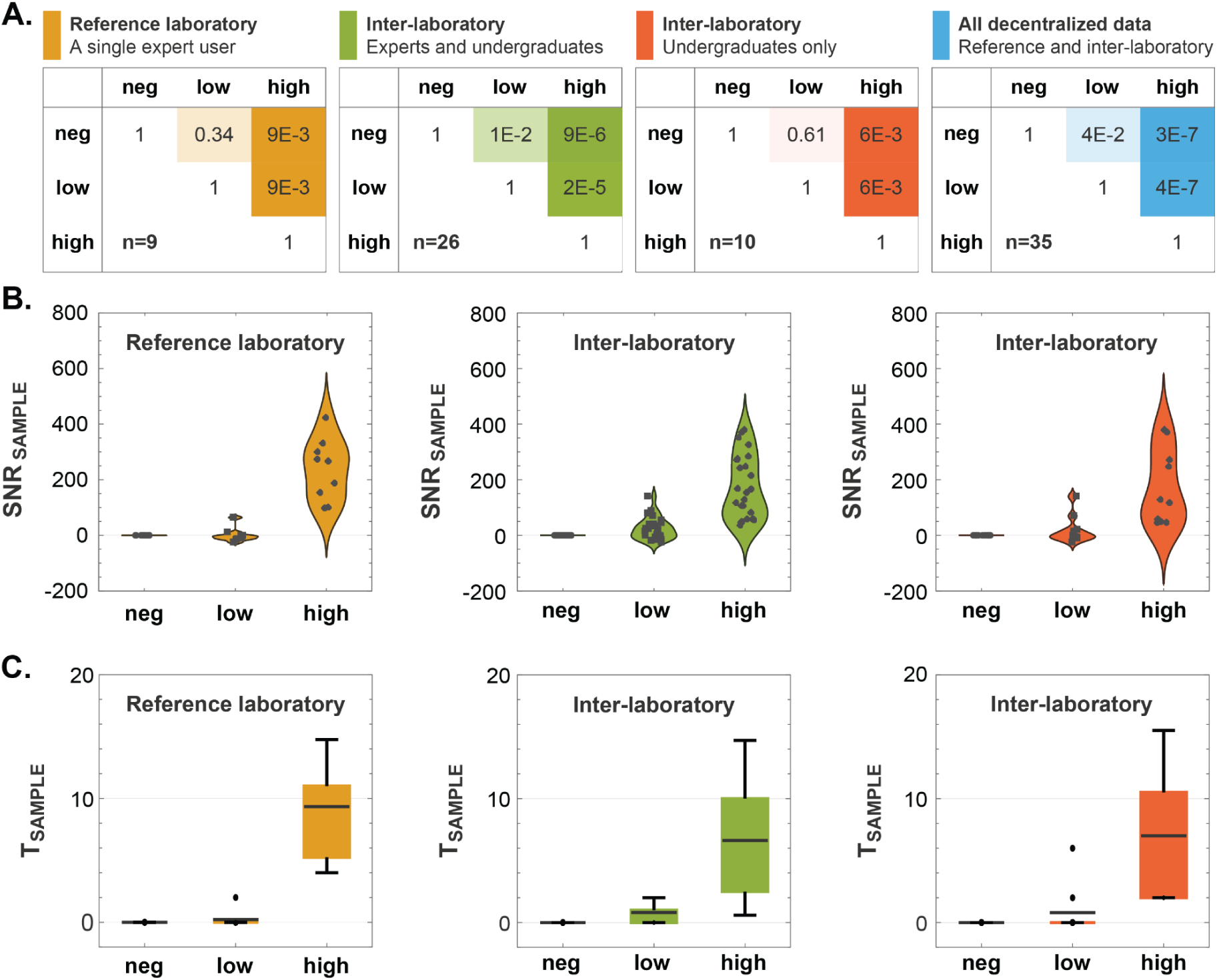
Discriminatory power of biocytometry across different segments of the decentralized data collection. **A)** A confusion matrix analysis illustrating the discriminatory power of biocytometry across different user groups in the decentralized data collection. Mean SNR values were computed across different user groups and for different HUMO samples. Wilcoxon signed-rank tests were conducted between different HUMO samples in each user group to test the hypothesis of equal medians. The p-values are represented visually, with lower values indicating lower probability of equal medians. **B)** Sample SNR distributions illustrated as violin charts. Scatter plots of the actual data points were overlaid on each figure. **C)** Box-and-whisker plots present illustrating sample target estimate distributions. A box-and-whisker plot is displayed for each HUMO sample type and segment of the decentralized collection with means, quartiles, and 5% and 95% quantiles.

Collectively, these p-values emphasize a clear distinction in the differentiation of the high HUMO samples. The subtler difference observed between the negative and low HUMO samples can be attributed to two things. Firstly, the HUMO samples, comprising temperature-sensitive human cell lines, may have experienced stress during transit, as suggested by the temperature monitoring detailed in S4 Figure. Bioparticles and other reagents within the biocytometry kits are robust to shipping conditions (see S5 Figure). Secondly, the limited number of assays within each user group, notably within the reference laboratory (n=9 assays) and among the undergraduate cohort (n=10 assays), could contribute to less pronounced differentiation. Intriguingly, the comparative analysis reveals that the proficiency level of the user does not alter the outcomes. Data from the broader spectrum of external participants, including both expert and undergraduate users (n=26 assays), achieved statistically significant results. This pattern persists across the entire set of decentralized data (n=35 assays), with the Wilcoxon signed-rank test yielding p-values of 4×10^−2^, 3×10^−7^, and 4×10^−7^ for comparisons between negative to low, negative to high, and low to high HUMO samples, respectively. These findings highlight the robustness of the biocytometry protocol and its capacity for reliable discrimination of HUMO samples, asserting that the technology is accessible and yields consistent results, even when applied by first-time users with less laboratory experience.

The consistent performance across various laboratory settings was further emphasized by the sample SNR distributions across individual user groups (Fig 5B). For the negative HUMO samples, the average SNR_SAMPLE_ was recorded at 0.0, with a maximum absolute deviation from this average being 0.2. This tightly bounded range of SNR values underscores the high specificity of the biocytometry measurements, confirming the expected absence of target cells. For the low HUMO analysis, average SNR_SAMPLE_ values were 0.7 (reference lab), 23.7 (inter-laboratory), 19.0 (undergraduates), and 16.6 (all decentralized data). For the high HUMO samples, the mean SNR_SAMPLE_ values reported were 237.1 (reference lab), 168.9 (inter-laboratory), 171.2 (undergraduates), and 186.4 (all decentralized data). Distribution of target cells within samples is in agreement with Poisson sampling distribution. Despite variations in laboratories, instruments, and user proficiency, the sample SNR distributions remain distinct, ensuring reliable target predictions across diverse settings.

Fig 5C illustrates the sample target estimate distributions. For the negative HUMO, target estimates across all user groups consistently registered at zero, with no instances of false positive readout. In low HUMO samples, an overall average of 0.7 target cells was observed. This estimate is well aligned with the expected 10^-5^ sensitivity equivalent (see Fig 1B). In high HUMO samples, the elevated SNR values yielded an average of 7.2 target cells, which is congruent with the 10^-4^ sensitivity equivalent, albeit slightly lower due to factors previously discussed related to sample stress during transit.

## Discussion

The promise of decentralized data collection in biomedical research and diagnostics hinges on the consistent and reproducible performance of technologies across diverse environments and user proficiency levels. Our comprehensive characterization of biocytometry, structured in three stages, offers significant insights and potential solutions for overcoming the challenges associated with decentralizing data collection in single-cell research.

First, we identified the target-specific conversion factor CF and validated the sensitivity, accuracy, and reproducibility of biocytometry in a centralized and controlled setting. The results demonstrate the ability of biocytometry to effectively discriminate between different HUMO samples (Wilcoxon signed-rank test, p-values 3×10^−5^, 1×10^−8^, and 1×10^−8^ when comparing negative to low, negative to high, and low to high HUMO, respectively). The agreement of biocytometry results with the absolute counts of target cells obtained by fluorescence microscopy highlights the ability of biocytometry to enumerate target cells. This stage underscored the inherent accuracy, precision and reliability of biocytometry, setting a solid foundation for subsequent evaluations.

In the second stage, we assessed inter-user reproducibility, with 51 assays conducted by multiple users in a centralized location using shared instruments. The results showed consistent SNRT values across users of varying proficiency levels. Mean well SNRT values of 18.9 (CV = 48.7%), 18.3 (CV = 60.6%), 19.6 (CV = 53.1%), and 19.2 (CV = 52.6%) were observed for routine, regular and first-time users, and all users combined respectively. The consistency was additionally supported by comparative analysis of target estimates in high HUMO samples (mean sample target predictions 16.8, 19.0, 20.0 for the three different users and 18.9 for all users combined). The technology’s inherent normalization against negative controls and its ability to directly enumerate target cells align with good clinical laboratory practice (GCLP) principles. This design ensures reproducibility and reliability of results, which are not significantly influenced by user variability across different operators.

The third stage of the study involved decentralized data collection, enlisting external participants to conduct assays in various laboratory settings. This phase was instrumental in validating the technology’s consistency and reliability in real-world, decentralized conditions. We showed that biocytometry can produce results of consistent quality across 12 different laboratories in spite of the varying equipment and conditions. This reproducibility was further highlighted by the diverse group of 22 users involved in the decentralized data collection, spanning from experienced proficient users to undergraduate first-time users. Such reproducibility implies a relatively flat learning curve, making the technology user-friendly and accessible from the outset, ensuring that GCLP standards are easily upheld even in the most decentralized settings.

Biocytometry performed equally well in decentralized settings and was robust to sample perturbation accrued during shipping. The HUMO samples shipped during decentralized studies were subject to fluctuations in temperature exceeding -30°C during one stage of shipment. Despite these perturbations, biocytometry effectively differentiated between the three types of HUMO samples (Wilcoxon signed-rank test, p-values 4×10^−2^, 3×10^−7^, and 4×10^−7^ when comparing negative to low, negative to high, and low to high HUMO samples, respectively). This performance is in contrast to the sensitivity of antibody-based methods to similar perturbations (data not shown). Our findings confirmed that the stability of the biocytometry kits themselves remained unaffected, suggesting that the inherent design and functioning of the technology provide a buffer against potential variables introduced by varying shipping conditions, different laboratory environments, or user proficiencies. The results from this stage, including the distinct distributions of the sample SNR values, further affirmed the robustness and applicability of biocytometry in diverse settings.

Biocytometry provides reproducible cell enumeration with a high level of sensitivity. This is achieved through two key features. First, a strong signal from activated bioparticles attached to target cells. Second, a zero false positive rate. Traditional methods like flow cytometry often struggle with high levels of noise, making it difficult to differentiate between samples with no targets and those with a single target [14]. On the other hand, biocytometry distinguished between negative, low, and high HUMO samples, even in decentralized data collection settings, and consistently produced zero target estimates for the negative HUMO samples across all user groups, as demonstrated by the narrow SNR range of -0.2 to +0.2 in the 78 assays. This precision highlights biocytometry’s capability to detect even the lowest concentrations of target cells accurately, without interference from false positives.

With biocytometry, our findings show that reproducibility does not necessitate a trade-off with sensitivity. This robustness is attributed to its foundational principle of biological identification of cell-specific immunological features in suspension. By prioritizing biology as the core driver, it minimizes the dependence on potential confounders and provides an accurate reflection of biological samples. Beyond single-cell research, the promising outcomes advocate for the broader adoption of decentralized data collection, potentially ushering in a new era of collaborative, multi-institutional research efforts. This shift could foster diversity of perspectives, enhance innovation, and lead to more comprehensive findings by involving a wider group of researchers. In summary, this study underscores the promise of decentralized data collection and highlights the benefits it can bring to the broader scientific community.

## Materials and methods

### Preparation of HUMO samples

#### Target cell line preparation

A HaCaT (keratinocyte) EpCAM+ cell line was obtained as a gift from research partners. Cells were grown in the recommended cultivation medium, DMEM + 10% FBS + Antibiotic/antimycotic mix, at 37°C, 5% CO_2_, regularly checked for mycoplasma contamination during their growth and discarded in case of positive results.

Before harvest, the cell concentration, confluency and morphology were inspected under microscope, and cells were labeled with a fluorescent label in order to facilitate differentiation of target cells from background cells for establishing precise target cell counts using microscopy. Cells were collected using the StableCell™ solution according to the manufacturer protocol, washed with fresh cultivation medium and treated with DNAse I for 5 minutes at 37°C. The suspension was filtered using a 20 μm filter, diluted to 2.5 M/ml concentration and incubated with ATTO425-Maleimide at 5 μM concentration for 15 minutes in the dark. Resulting fluorescently tagged cells were washed with PBS + 0.1% gelatin twice and resuspended in the cultivation medium at 1 M/ml concentration. Cell concentration was established with a Burker chamber.

#### Background cell line preparation

An HL-60 (promyelocytic) EpCAM- cell line was obtained as a gift from research partners. Cells were grown in the recommended cultivation medium, RPMI + 10% FBS + Antibiotic/antimycotic mix, at 37°C, 5% CO_2_, regularly checked for mycoplasma contamination during their growth and discarded in case of positive results.

Before harvest, the cell concentration and morphology were inspected under a microscope. Cells were harvested by spinning down 10 ml of the cell culture at 250 RCF for 5 minutes. The supernatant was discarded and the pellet was resuspended in 1 ml of RPMI++ medium. The suspension was filtered through a 10 μm filter. Cell concentration was established with a Burker chamber.

#### Preparation of the HUMO samples

The volume necessary to obtain 10^6^ of HL-60 cells was calculated and transferred to a fresh microcentrifuge tube. No spiking was done for the negative HUMO sample. The low HUMO sample received a spike of 250 HaCaT cells. The high HUMO sample was spiked with 4,100 HaCaT cells. Each tube was then filled up to 500 μl with RPMI++ medium.

For quality control, both the cell count and viability were established. A 10 μl sample of the cell suspension was combined with 10 μl of Trypan Blue and incubated at room temperature for 5 minutes. The cell count was ascertained using a Burker chamber and the number of dead cells was recorded. In addition, the concentration of target cells was identified through fluorescence microscopy. Five aliquots of 10 μl each were transferred to a 96-well plate to determine the concentration of fluorescent targets.

Next, 400 μl of the cell suspension was transferred to a fresh tube, and 400 μl of a mixture of 80% iFBS and 20% DMSO was added. This mixture was then gently vortexed and aliquoted by 10 μl into the final tubes and capped. Sample tubes were moved to Mr. Frosty containers and stored at -80°C. The tubes were transferred to their designated storage boxes at -80°C after a 3-hour incubation at -80°C.

### Biocytometry kit

Every participant was provided with a biocytometry kit (Sampling Human, Berkeley, CA, USA) comprising 14 tubes and accessories:

● **Resuspension Buffer (1x)** - Resuspension buffer is used to dilute the HUMO samples and provide an appropriate environment for the reaction.
● **HUMO Samples (3x)** - HUMO samples containing EpCAM+ HaCaT target cells integrated into a leukocyte surrogate population of 10^6^ HL-60 EpCAM- cells. Samples were preserved in a cryopreservation medium. Each biocytometry kit contains one tube of each sample type characterized by different target cell concentrations - negative (absent of target cells), low (10^-5^ sensitivity equivalent), and high (10^-4^ sensitivity equivalent).
● **Reaction Tube (3x)** - Engineered bioparticles, the active component of the assays, supplied in a desiccated form and rehydrated upon addition of the sample.
● **Master Tube (3x)** - Each master tube containing 1000 μL of proprietary medium, a semi-permeable nutritive medium with thermoresponsive properties in which the sample processing takes place.
● **Luminescence Buffer (3x) and Luminescence Substrate (1x)** - Luminescence buffer and substrate are mixed for readout of processed samples. Composition of the luminescence buffer supports strong luminescence signals and ensures its stability for up to 30 minutes at room temperature. Each luminescence buffer tube contains 975 µl of reagent. The luminescence substrate tube contains 200 µl of substrate resuspended in ethanol.
● **Evaluation Strip (3x) and Plate Adapter (1x)** - A combination of three 8-tube PCR strips and a PCR strip rack is used for the luminescence readout in a microplate reader. Biocytometry kits were shipped to participants on dry ice and stored at -80°C upon reception until their use.

### Biocytometry assay protocol

Every participant was provided with a comprehensive protocol in the form of a PDF document (see S1 File). This protocol outlined detailed instructions encompassing the handling of the biocytometry kit, sample preparations, step-by-step procedural guidance, basic configurations for luminescence readout, and troubleshooting tips. Furthermore, to enhance clarity and offer a visual guide, participants were advised to watch a video tutorial detailing the same instructions (see S2 Movie). The subsequent sections provide a simplified overview of the biocytometry assay protocol.

The biocytometry assay commences with the addition of 90 µl of **Resuspension Buffer** to each **HUMO Sample**. The entire volume (100 µL) from each **HUMO Sample** is then transferred to an individual **Reaction Tube** without any mixing and is incubated for 5 minutes. The contents of the **Reaction Tubes** are resuspended by gently pipetting up and down 10 times.

The **Reaction Tubes** are centrifuged with the notch oriented away from the center at 200 RFC for 1 minute. Each **Reaction Tube** is then twisted 180° until the notch faces inward, and centrifuged a total of 10 times, rotating the tube 180° between each spin down. The contents of the **Reaction Tubes** are then gently resuspended by pipetting up and down 10 times.

75 µl from each **Reaction Tube** is transferred to each **Master Tube**, and mixed by inverting 10 times. The **Master Tubes** are then incubated at 37°C for 5 minutes, and subsequently mixed by inverting 10 times. One **Evaluation Strip** is filled from each **Master Tube** by dispensing 75 µl into each well. The **Evaluation Strips** are then incubated in a thermocycler using a predefined program consisting of four cycles. First, a setting phase at 4°C for a duration of 10 minutes. Second, a processing phase at 30°C for 4 hours. Third, a deactivation step at 50°C for 15 minutes. Fourth, the sample can be held at 4°C for an extended period of up to 24 hours if delayed readout is needed (optional). The thermocycler lid temperature is set to 35°C for all steps.

The plate reader is adjusted to the recommended settings (luminescence readout from top, high gain, no attenuation, probe as close to the labware as possible). **Readout Reagent** is prepared by adding 25 µl of **Luminescence Substrate** to each **Luminescence Buffer** tube and mixed by vortexing. The **Readout Reagent** (75 µl) is added to each well of the three **Evaluation Strips**. The **Evaluation Strips** are incubated at 37°C for 5 minutes, placed into the **Plate Adapter**, and then incubated upside down at room temperature for 1 minute. The **Evaluation Strips** are centrifuged at 1000 RCF for 1 minute. Measurements are then taken using the microplate reader and the **Plate Adapter** according to the recommended settings. The results are reported online.

### Readout & data collection

Data collection of biocytometry assays from the internal studies were collected with the following plate reader settings.

**Detection method:** Luminescence

**Optics type:** Luminescence fiber / filter

**Read height:** Default

**Labware:** 96-well plate with lid (standard ANSI/SLAS compliant 96-well microplate)

**Read from:** Top

**Gain:** 255 / no attenuation (decrease as necessary to prevent signal overflow)

**Read type:** Endpoint / kinetic

**Integration time:** 1 s

**Interval:** 1 minute or minimum possible

**Number of reads:** 5 (average used as the final readout values)

**Read area:** 4 columns (3 samples, 1 blank)

The same specifications were provided to the participants of the study with the understanding that individual adjustments would be made depending on the make and model of the instruments available.

Each assay measured the three provided samples as well as one blank column containing no sample and no evaluation strip. Users were asked to report the averaged values of the five kinetic reads to the online portal. In addition to the measured values, metadata regarding the plate reader model, read settings, and additional comments were collected for each assay performed. Participants were asked to self-report in the comments whether any steps were omitted or changed during the preparation of the assay. These notes were then used to discard any assays which did not adhere to the standard protocol.

### Assessment and validation of assay data integrity

Upon completion of data collection, assays from the decentralized phase were classified as valid or invalid, based solely on procedural adherence. No criteria rooted directly in the data itself were used to exclude any assays. This approach prioritized adherence to the established protocol over the outcomes observed in the collected data.

The assessment of assay validity involved a thorough two-step process to ensure rigorous protocol adherence. Initially, participants were asked to report any significant deviations or issues for each assay submitted. Subsequently, each participant was contacted by one of the authors to discuss any deviations from the protocol or issues encountered during assay preparation. If no gross protocol failures were identified during this discussion, the assay was deemed valid. Assays were considered valid unless accompanied by acknowledged deviations or preparation issues categorized as gross. This meticulous process was crucial in filtering out data potentially compromised by significant non-adherence to the protocol.

Following the data collection and evaluation process, several significant deviations from the protocol were identified, including:

1. Deviation from protocol, such as imprecise timing (2 assays)
2. Liquid handling that diverged from the specifications (13 assays)
3. Storage of the kit contrary to outlined instructions (7 assays)
4. Data acquisition using readers with no luminescence readout support or too low sensitivity (6 assays)
5. Readout issue or other instrument malfunction (10 assays)
6. Sample labeling and tracking issue (4 assays)

A total of 120 biocytometry kits were distributed for decentralized data collection: 14 to the reference lab and 106 to external participants. Of these, 71 assays were completed and their results submitted online. Only 35 assays met the validity criteria and were included in subsequent data analyses, while 36 were categorized as invalid due to the deviations listed and excluded from further consideration. The remaining kits were not used due to the unavailability of required instrumentation or lab personnel.

### Data normalization

For consistent data interpretation, normalization was an essential step. Every biocytometry assay incorporated a negative HUMO sample, which was designated as a negative control. These negative HUMO samples were devoid of target cells but contained an equivalent number of background cells, establishing them as an ideal reference to adjust for potential disparities that may arise due to variations in microplate readers, incubation conditions, sample handling, and other potential sources of variance.

The assay measurements captured raw luminescence data from each of the three sample types. Each sample was distributed across 8 wells, and each well was subjected to 5 kinetic reads. From this raw data, luminescence readings were averaged across the five measurement points, generating a value denoted as LUM_WELL_ for each well. Subsequently, only this average value was pursued in further analyses.

For each assay, the assay negative control denoted as *NC*, was computed by averaging the luminescence output over the 8 wells designated as the negative control. This NC, in conjunction with LUM_WELL_, was instrumental in calculating the signal-to-noise ratio of each well SNR_WELL_ within the assay, given 24 wells in total. Assuming noise to be 10% of the control value, the SNR for each well was computed as:

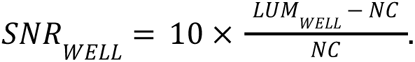

The resulting SNR values underwent various metrics evaluations considering both the individual SNR_WELL_ values, and the sample mean SNR denoted as SNR_SAMPLE_. The SNR_SAMPLE_ was deduced by summing up the SNR_WELL_ values across its 8 respective wells.

### Validation by microscopy and SNRT evaluation

To validate the number of target cells present in the reaction, the contents of the evaluation strips were transferred to a 96-well plate with flat clear bottom for observation by fluorescence microscopy. To do so, evaluation strips were heated to 37°C for 5 minutes in a thermocycler after luminescence readout by biocytometry. Using a multichannel pipette, the contents of the strips were pipetted up and down 3-5 times to resuspend the cells from the bottom of the tube, then transferred to a flat bottom 96-well plate. The contents of each single tube of 200 µL was transferred to a corresponding well in the plate. During this process, some liquid - and consequently, target cells - could be lost, either when removing strip lids or from residual volumes that could not be fully transferred. Additional target cell losses may occur if cells remain adhered to the strip walls or are left in any residual liquid.

The plate was heated again for 10 minutes at 37°C in a thermocycler and immediately transferred for centrifugation at 1000 RFC for 1 minute, bringing the cells to a single plane at the bottom of the plate. Each well was scanned in its entirety on an inverted microscope under a CFP fluorescence filter to count the number of fluorescently stained target cells. The number of target cells obtained by microscopy is denoted as T_WELL,MICROSCOPY_.

The T_WELL,MICROSCOPY_ values could then be validated against the corresponding luminescence measurement in each tube of the evaluation strip in order to establish the true signal-to-noise ratio per target (SNRT) value as follows:

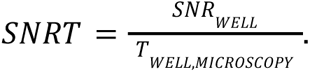

### Evaluation of the target and SNRT estimate

The number of target cells in each well, denoted as T_WELL_, is estimated by normalizing the SNR_WELL_ value by the conversion factor CF, which was identified during the benchmarking of biocytometry in combination with the microscopic analysis. The resulting value is rounded to the nearest integer, reflecting the physical reality that only whole cells can be present. Thus, T_WELL_ is calculated as:

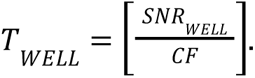

The total number of target cells in the analyzed sample, T_SAMPLE_, is then the sum of T_WELL_ values across its 8 respective wells.

The estimate of the SNRT value, denoted as SNRT_EST_, serves as an indicator of assay consistency. It validates the results of assays not accompanied by microscopy by ensuring congruence between biocytometry estimates and actual target cell counts. SNRT_EST_ for each well is calculated by normalizing the SNR_WELL_ value by T_WELL_ and rounding the result to the nearest integer, as follows:

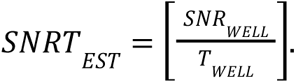

## Acknowledgements

Undergraduate faculty and student participation was supported by the Cell Biology Education Consortium: Path to Publication, an NSF-funded RCN-UBE (Award ID #2316122).

## Supporting information

**S1 File. Biocytometry assay protocol.** Detailed instructions were provided to all participants regarding biocytometry kit and samples handling, assay execution, luminescence readout configuration and troubleshooting tips.

**S2 Movie.** Biocytometry assay video protocol. Detailed video instructions were provided to all participants guiding them through the biocytometry assay execution.

**S3 Dataset.** Data collection spreadsheet. Spreadsheet containing data collected from internal users and external participants with raw luminescence readouts and data converted to SNR values and target cell estimates. Only technical entries for the data are included, and participant names have been replaced by numbers.

**S4 Figure. Shipment temperature.** The figure details temperature monitoring for the shipment of HUMO samples and biocytometry kits from the centralized laboratory to the reference decentralized laboratory. Shipped with the expectation of maintaining -80°C on dry ice, a significant deviation was recorded by the temperature monitor between hours 144 and 192, with a peak at around -27°C. Given the temperature sensitivity of the HaCaT and HL-60 cell lines contained in the HUMO samples, this unexpected rise in temperature may have compromised their viability, potentially leading to cellular necrosis and impacting the integrity of the samples upon arrival.

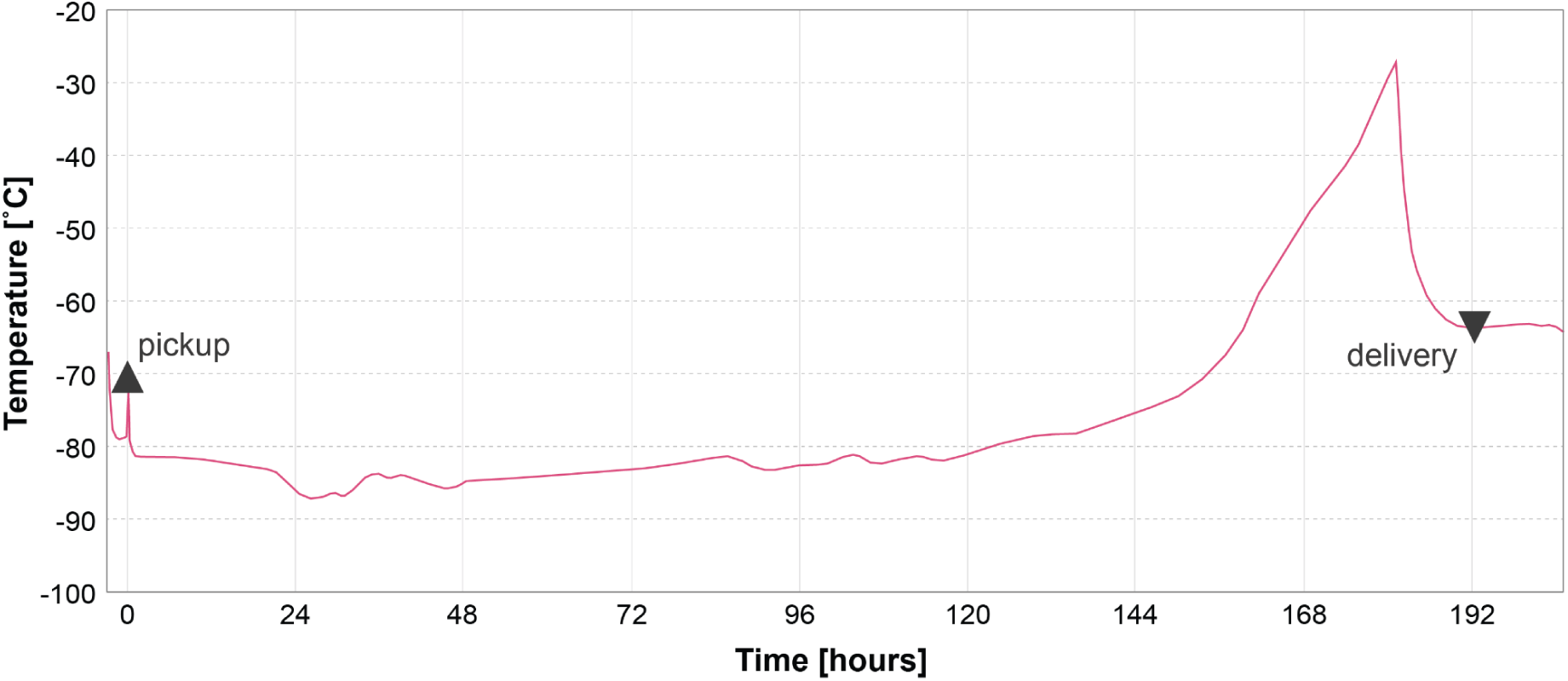

**S5 Figure. Stability of bioparticles.** A longitudinal stability study of the bioparticles integral to the biocytometry kits demonstrated sustained performance, with no significant decline observed over a 16-week period when stored at -20°C, as depicted in S5 Fig. This stability indicates the temperature fluctuations recorded during the shipment, detailed in S4 Fig, did not compromise the efficacy of the biocytometry kits. Consequently, these findings validate the robustness of the study design and assure that the assay results remained unaffected by the temperature variances encountered in transit.

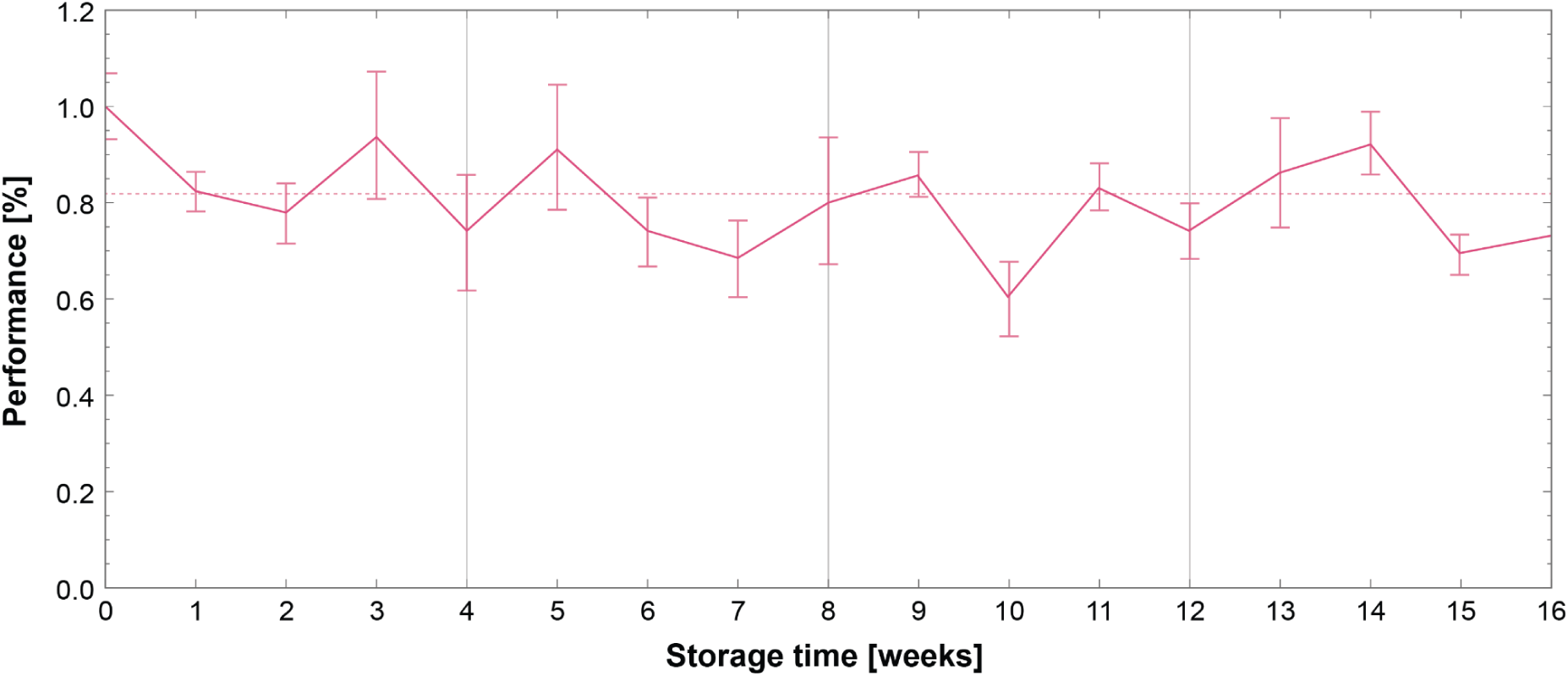

**S6 Supporting information.** De-hashing decentralized data. In order to ensure an unbiased analysis in the decentralized data collection phase, the HUMO samples were delivered to participants under a disguised nomenclature, obscuring the identity of each sample. The samples were labeled as HUMO A, HUMO B, and HUMO C instead of their actual types; low, high, and negative, respectively. Participants, therefore, engaged in blind analysis, unaware of the specific HUMO type they were analyzing. Upon receiving the data from the participants, a critical step of data de-hashing was performed. This involved reorganizing the submitted data to reflect the true identity of the samples, aligning them to their respective types - negative, low, and high HUMO. Through this meticulous process, the data was aptly restructured for subsequent analysis, while assuring an unbiased data collection and validation process.

**S7 Supporting information.** Box charts of target estimates. Sample target estimates were split based on the respective sample type and assay cohort. Sample target estimates were then visualized in Figs 3C, 4B and 5C using a box chart for different users and assay cohorts. In the box chart, each box consists of a median marker, fences that represent quartiles, and whiskers that represent 5% and 95% quantiles. Quantiles were estimated using a mode method. Outliers were detected by a median based method. Individual well target estimates were visualized and compared to the target cell counts obtained by microscopy in Fig 2C.

**S8 Supporting information.** Hypothesis testing. A hypothesis test was used to judge the ability of the technology to discriminate between sample types as follows. Assays were categorized into distinct cohorts based on assay and user metadata (e.g. distinct laboratory or user proficiency level). For each sample type, the sample means were consolidated across assays within a specific cohort. A Wilcoxon signed-rank test was subsequently executed, comparing the sample means of type i with those of type j within the given assay cohort. The p-value for this test was determined for every pertinent assay cohort and for each pairing of sample types. The outcomes of these tests were graphically represented in Figs 3A and 5A.

**S9 Supporting information.** Histograms. SNRT values were further visualized using a smooth histogram in Fig 2B. The histogram was smoothed with a gaussian kernel.

