## Supplementary figures and images for "Enhancing reproducibility and decentralization in single cell research with biocytometry"

### Supporting Figure 4

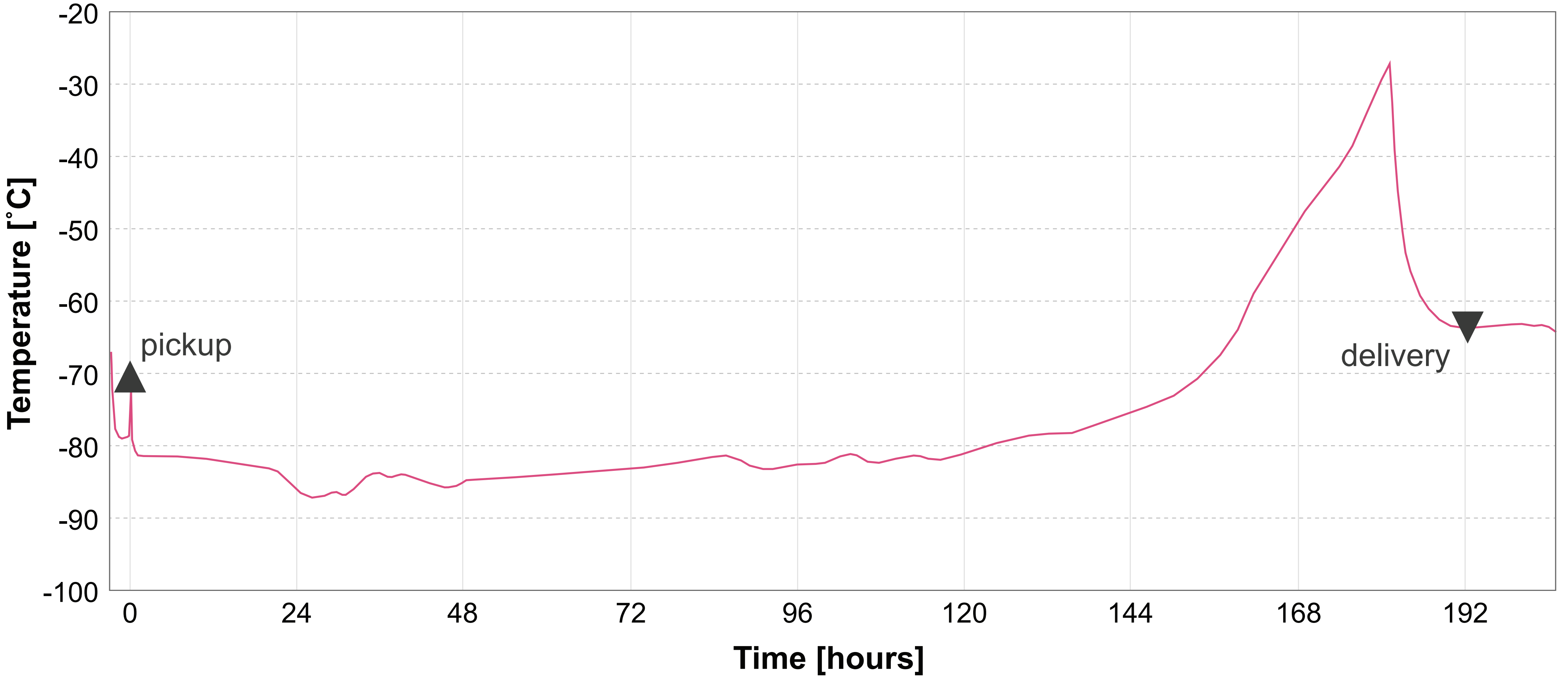

### Supporting Figure 5

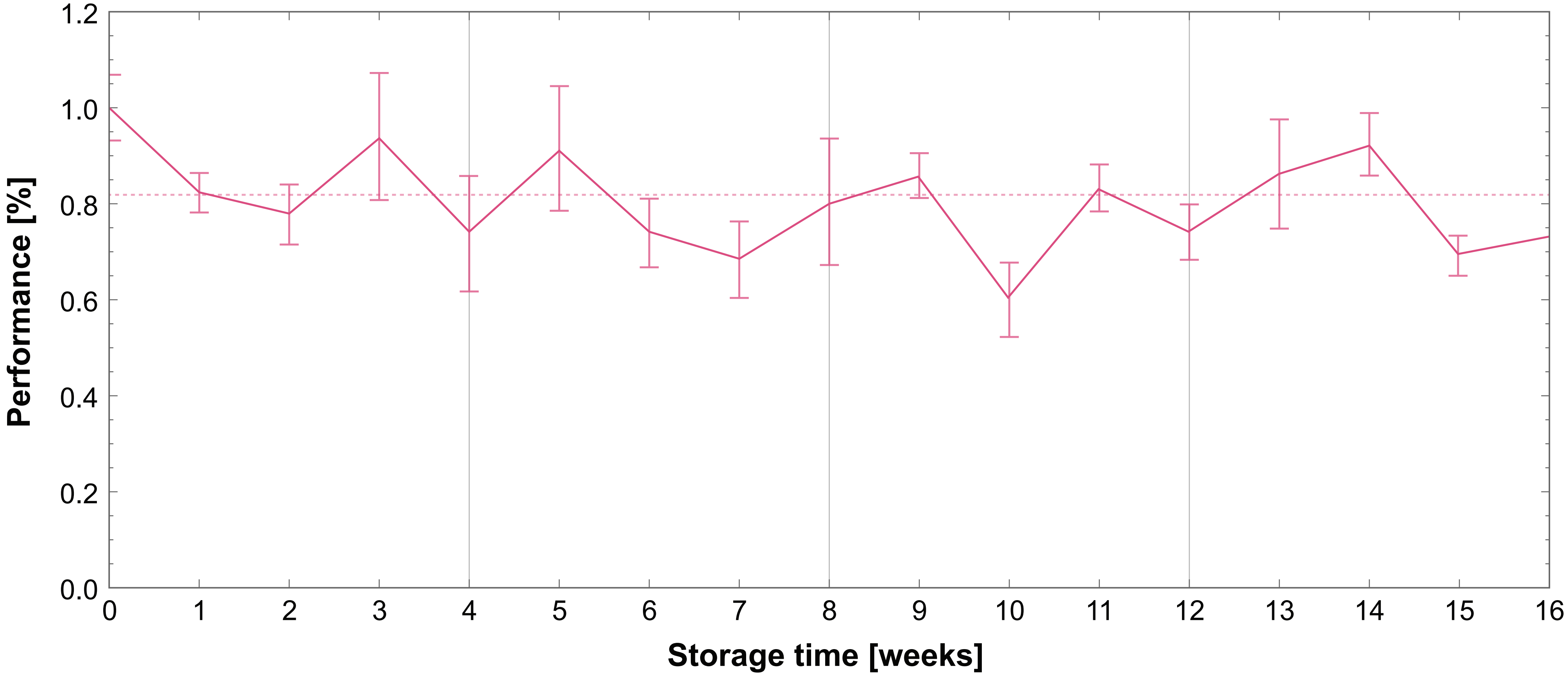
