## Supporting File 1 for "Enhancing reproducibility and decentralization in single cell research with biocytometry"

**DOT Assay Full Protocol - (Version 2.2, updated May 9, 2023)**

The **HUMO Samples** have been labeled as A, B, and C, but participants will not know which phenotype this corresponds to. Only after measurement will users know which HUMO is which. It is important to always keep track of which sample is A, B, and C, so that the matching phenotypes can be later validated.

⚠ *Please keep in mind that the **HUMO Samples** contain mammalian cells and should therefore always be handled gently, and always mixed before transferring to prevent settling!*

### Unpacking

1. Record the **Kit ID** in your lab notebook or by taking a photo. This will be used when reporting your results.
2. Remove one **DOT Run** package and return remaining DOT kit contents to -80°C.
3. Remove the packet of **Temperature Sensitive Reagents** (three **HUMO Samples** and one **Luminescence Substrate**) and place them at -80°C until further use.
4. The remaining contents of the **DOT Run** package can be kept at room temperature until their use.

### Preparation

5. Preheat a water bath or heat block to 37°C.
6. Place **Master Tubes** (blue) and **Resuspension Buffer** (white) at 37°C.
7. Transfer the **Resuspension Buffer** to room temperature once fully thawed (3-5 minutes). Vortex to ensure that no ice remains in the lid of the tube.
8. Leave the **Master Tubes** at 37°C until later use.
9. Centrifuge the three **Reaction Tubes** (black) at 200 × *g* for 30 seconds to collect the dried reagent at the bottom of the tubes.
10. Label one of each **Reaction Tube**, **Master Tube**, and **Evaluation Strip** as A, B, and C. Do not write on the lid of the **Evaluation Strip** tubes as it will interfere with the readout, use the upper tab instead.

#### DOT Mixing

11. Transfer **HUMO Samples** (red) from -80°C storage directly to 37°C for 3 minutes.

    ⚠ *The HUMO Samples are of low volume and thaw quickly, only transfer to 37°C when you are full prepared and ready to begin.*
12. Centrifuge the **HUMO Samples** at 50 x *g* for 30 seconds to collect contents.
13. Add 90 µl of **Resuspension Buffer** to each **HUMO Sample**.
14. Mix the **HUMO Samples** by **gently** pipetting up and down before transferring 100 µl (entire contents) of each **HUMO Sample** to its respective **Reaction Tube.** Pipette directly onto dried reagent to ensure proper rehydration. Do not mix after.

    ⚠ *Pipette up and down immediately before transferring to prevent loss of targets*
15. Let the reagent rehydrate for 5 minutes at room temperature.
16. Resuspend the three **Reaction Tubes** by pipetting up and down 10 times gently or until the solution is homogeneous.  Avoid creating bubbles.
17. Place the **Reaction Tubes** into a fixed-angle centrifuge, orienting the notch away from the center, and spin down at 200 × *g* for 1 minute.
18. Without removing the tubes from the centrifuge, twist the **Reaction Tubes** 180° until the notch faces inwards to the center (see Figure 1).
19. Spin down 9 more times in the same manner, rotating the tube 180° between each spin down.  A small pellet should be visible.
20. Resuspend the three **Reaction Tubes** by pipetting up and down 10 times gently or until the solution is homogeneous with no visible pellet.  Avoid creating bubbles.

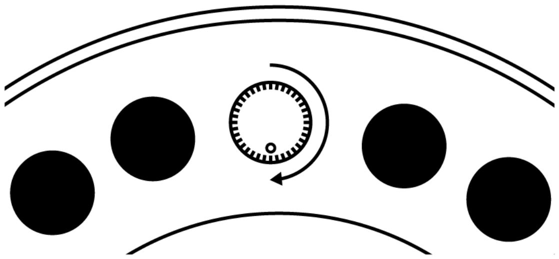

Figure 1.

**DOT Incubation**

21. Retrieve the three **Evaluation Strips** and place them into a **Plate Adapter**.
22. Retrieve the three **Master Tubes** from the 37°C.
23. Transfer 75 µl from each **Reaction Tube** into the corresponding **Master Tube** and mix by inverting 10 times.

    ⚠ *The gel medium should be fully liquid and should move when the tube is inverted*

24. Incubate the **Master Tubes** at 37°C for 5 minutes.
25. Retrieve the **Master Tubes** and invert 10 times.
26. Dispense 75 µl from each **Master Tube** into each well of the corresponding **Evaluation Strip** by reverse pipetting. Do not change tips between wells.

    ⚠ *The medium will solidify at room temperature in 6-10 minutes, so it is important to work quickly.*

27. Place the lids on the **Evaluation Strips** and press down firmly.
28. Run the **Evaluation Strips** in a thermocycler using the following program:

| Step | Temperature | Time |
|---|---|---|
| Setting | 4° C | 10 minutes |
| Processing | 30° C | 4 hours |
| Deactivation | 50° C | 15 minutes |
| Hold | 4° C | up to 24 hours |

*The lid temperature is set to 35° C for all steps, no more than 40°C.

**DOT Measurement**

29. Retrieve **Luminescence Substrate** (yellow) from -80°C storage and three tubes with **Luminescence Buffer** (green) from room temperature storage.
30. Prepare the readout reagent by adding 25 µl of **Luminescence Substrate** to each of the three **Luminescence Buffer** tubes. Vortex the tubes thoroughly to ensure complete mixing.
31. Add 75 µl of the **Luminescence Buffer** with added substrate to each well of the three **Evaluation Strips**.
32. Press the lids of the **Evaluation Strips** firmly into place.
33. Place all three **Evaluation Strips** in the thermocycler and incubate for 5 minutes at 37°C to melt the contents.

    ⚠ *Do not incubate for longer than 5 minutes as this may compromise the signal*
34. Firmly push the **Evaluation Strips** into the **Plate Adapter** and place them lid side down at room temperature for 1 minute to allow for mixing, the gel will not move.
35. Place the **Plate Adapter** in an upright position and centrifuge 1000 × g for 1 minute.
36. Check that the **Evaluation Strips** are snapped firmly into place in the **Plate Adapter** and that all the lids are pushed down as far as possible.
37. Measure the **Evaluation Strips** in a plate reader, using the **Plate Adapter** and the following settings (see page 10 for model specific settings):

    Detection Method: Luminescence

    Optics Type: Luminescence Fiber/Filter

    Read Height: Default

    Plate Type: 96-well plate **(LID ON)**

    Read from: Top

    Gain: 255 (decrease as necessary)

    Read Type: Endpoint/Kinetic

    Integration Time: 1 s

    Interval: 1 minute or minimum possible

    Number of reads: 5 (average used as the final readout values)

    Read area: 4 columns (3 samples, one empty)

### Reporting Results

38. Log in using your **Kit ID** at www.samplinghuman.com/r-prj
39. In the upper right menu, click on My Kit to open the data submission form.
40. Data can be directly copy-pasted from an excel sheet into the web form.
41. Enter the additional metadata regarding your assay.

   ⚠ *If any deviations from the protocol occurred, record them in the comments section.*

42. Click "Save Run" to send your data. Once you have submitted your data, you will not be able to edit it again. If your data has been submitted in error, please contact
43. Give us a rating and provide feedback on your experience in the follow-up survey.
44. Perform another DOT or check out updates on the project on the website.

**Thank you for participating in the Reproducibility Project!**

### Troubleshooting Guide

#### I. Unpacking

| Observation | Action |
|---|---|
| DOT kit was not stored at -80°C.<br>(Step 2) | The HUMO samples contain mammalian cell lines that are particularly sensitive to changes in temperature. If your kit was stored at above -80°C for any extended period, the samples are compromised and cannot be included in the published data. |

#### II. Preparation

| Observation | Action |
|---|---|
| Reaction Tube looks empty.<br>(Step 9) | The desiccated reagent is clear and often hard to see. If the solution does not become cloudy during rehydration, or there is no pellet visible after the centrifugation steps, there may be a manufacturing error. This replicate will have to be discarded. |
| Lids of Reaction Tubes were not fully screwed on in the package.<br>(Step 9) | The desiccated reagent is vacuum packed to ensure that it does not come in contact with air or moisture. The lids are loosely attached to ensure that air within the tubes is removed as well. You can proceed as normal with the assay. |

#### III. DOT Mixing

| Observation | Action |
|---|---|
| Centrifuge cannot reach speeds as low as 50 x *g*.<br>(Step 12) | Set your centrifuge to the lowest speed possible. Record the actual speed in the notes when you submit your assay. |
| Volume transferred from resuspended HUMO Sample was significantly less than 100µL.<br>(Step 14) | Spin down the HUMO sample at 50 x g for 30 seconds before adding the resuspension buffer. If the sample was stored for an extended period, some loss may have occurred due to evaporation. Proceed with the assay as usual. |
| Cannot resuspend dried reagent.<br>(Step 16) | If the dried reagent does not easily become resuspended after 5 minutes of rehydration, the Reaction Tube may have become compromised. The reagent may appear white before |

| | |
|---|---|
| | rehydration in these cases. This replicate will have to be discarded. |
| Cannot resuspend pellet after centrifugation. (Step 20) | Check that you have used the correct settings for centrifugation (200 x g, 1 min each). The pellet should resuspend easily after pipetting up and down 10 times. Replicates in which the pellet could not be resuspended will have to be discarded. |

### IV. Incubation

| **Observation** | **Action** |
|---|---|
| The Master Tube is full of small bubbles. (Step 22) | This is normal to see when the medium has been frozen. It will not affect the readout, proceed as usual. |
| The Master Tube medium does not move when inverted. (Step 23) | The contents of the Master Tube should be fully liquid when you are working with it. Check that it has reached a temperature of 37°C and that it has been preheated for at least 10 minutes before handling. Do not transfer the reaction to the Master Tube unless the medium is in a liquid state. |
| The Master Tube medium is too viscous to dispense accurately. (Step 26) | Depending on the ambient temperature, after being removed from 37°C the medium has roughly 6-10 minutes of working time before it becomes too viscous to handle. If you find that there is not enough time to dispense all three Master Tubes, you may consider holding the remaining tubes in polystyrene as you dispense to keep them from cooling too quickly. If it is necessary to return the Master Tube to 37°C to re-melt the contents for dispensing, please note that you have done so when reporting your results. |
| The gel medium is not distributed equally in the wells of the evaluation strip. (Step 26) | Dispense the gel medium by reverse pipetting. The medium is viscous as it cools, so it is slower to fully aspirate. Allow the tip to completely fill before removing it from the medium to avoid bubbles in the tip of the pipette. |

| | |
|---|---|
| There is a bubble trapped at the bottom of the evaluation strip after dispensing medium.<br>(Step 26) | A small bubble is normal, and will not affect the readout in any way. However, it is best to avoid large bubbles as it may take up space necessary for the readout buffer. To avoid this, dispense the medium as close to the bottom of the well as possible without touching the tip directly. |
| Volume exceeds recommended volume of thermocycler.<br>(Step 28) | This should not cause any problems provided that the lid of the thermocycler is on tight. |
| Cannot achieve the recommended lid temperature.<br>(Step 28) | If your thermocycler cannot reach 35°C, you may use a temperature up to 40°C, but not above. If that is not possible, switch off lid heating completely and note it in the comments when reporting your results. |
| Plate was held at 4°C before readout for longer than 24 hours.<br>(Step 28) | This may result in high signal levels overall. Please make note of roughly how long the plate was incubated when reporting your results. |
| Condensation present on the lids of strips after incubation.<br>(Step 28) | A small amount of condensation of the lids of the strips is not unusual after incubation. If large droplets are present, allow the gel medium to cool to a fully solid state before centrifuging (30 seconds, 1000 x *g*). Make note of this extra centrifugation step if it was performed. |

#### V. DOT Readout

| **Observation** | **Action** |
|---|---|
| The medium does not move when the Evaluation Strips are inverted.<br>(Step 34) | It is expected that the medium will not fall to the bottom of the strips due to its viscous nature. The medium will mix with the readout reagent regardless, as long as the strips are inverted immediately after being removed from 37°C. |
| Plate too tall for plate reader.<br>(Step 37) | Check that the Evaluation Strips are snapped into place and the lids are completely flat and level. Make sure that you have selected the setting to measure with the lid on. |
| Signal overflow.<br>(Step 37) | Decrease the gain in increments of 50 until there is no longer overflow. Make note of the gain settings when reporting your results. |
| Low signal overall.<br>(Step 37) | Check that the gain is set to 255. Reasons for low overall signal include: |

| | |
|---|---|
| | - Luminescence Substrate degradation: Store substrate at -20°C or below and do not expose to light.<br>- Reaction Tubes compromised: Unexpected temperature fluctuations or freeze-thaw may compromise the DOTs.<br>- Luciferase degradation: Do not allow the plate to sit for an extended period after adding readout buffer. Extended heating or excess temperatures may adversely affect signal levels.<br>- Individual differences between plate readers. Low signal values are not necessarily indicative of a failed assay. |
| Low signal to noise ratio<br>(Step 37) | If all steps of the protocol were performed without issue, it is possible that the HUMO samples were compromised from temperature fluctuations or improper handling. Please report your results anyway and contact for further assistance. |
| Inconsistent signal values within a single sample.<br>(Step 37) | It is expected that significant variation will occur in the high HUMO phenotype due to the random distribution of targets. If you have any questions about how to interpret your results, you can contact for further assistance. |

### Reader Specific Settings

R.PRJ DOT kits are supplied with a set of the ThermoFisher Scientific VersiPlate labware. The labware dimensions are identical to standard ANSI/SLAS compliant 96-well microplates.

When setting up your reader, use standard 96-well plate labware definition with lid. If configuring the plate height manually, consider the total height including lids to be 17 mm. The optimal read height allows reader optics to be as close to the plate as possible, while avoiding well-to-well signal crosstalk.

The following table shows configurations for some of the most common microplate readers.

| Manufacturer | Model | Settings |
| --- | --- | --- |
| Biotek | Synergy H1<br>Synergy HT | • Labware: 96 WELL PLATE<br>☑ Use lid<br>• Integration time: 1 second<br>• Read height: 4.5 mm<br>• Gain: 255 (lower to 200 if values overflow and repeat) |
| Biotek | FLx800 | • Labware: 96 WELL PLATE<br>☑ Use lid<br>• Integration time: 1 second<br>• Gain: 255 (lower to 200 if values overflow and repeat) |
| Biotek | Cytation 1 | • Labware: 96 WELL PLATE<br>☑ Use lid<br>• Integration time: 1 second<br>• Read height: 12 mm<br>• Gain: 255 (lower to 200 if values overflow and repeat) |
| Tecan | Spark 20M | • Labware: [NUN96fw_Lumi_Nunc Fluoror Nunc] - ThermoFisher Scientific-Nunclon 96 Flat white<br>☑ Plate with cover<br>• Attenuation: None<br>• Integration time: 1000 ms (lower if needed and repeat)<br>• Settle time: 0 ms |
